# The gene *yellow* influences aversive mate preference learning in a butterfly

**DOI:** 10.64898/2026.07.10.737617

**Authors:** Yi Ting Ter, David A. Ernst, Keity J. Farfán-Pira, Erica L. Westerman

**Author notes:** Corresponding author – 850 W Dickson St, Fayetteville, AR 72701, USA, E. L. Westerman.

## Abstract

Reproductive isolation is a central driver of speciation and can be reinforced by both innate mate preferences and mate preference learning. Pigmentation genes are strong candidates for pleiotropic effects on these processes because they shape visual traits used in mate choice and may also influence neural function, yet their role in learning is poorly understood. Here, we test whether the pigmentation gene *yellow* affects innate mate preference and aversive mate preference learning in female *Bicyclus anynana*, and whether these effects are associated with changes in brain dopamine levels. Using CRISPR-Cas9, we knocked out the *yellow* gene, developed a mutant line, and tested the mate preferences and mate preference learning ability of mutant females compared to wild-type (WT) females. We find that loss of *yellow* does not alter assortative mating based on pigmentation or disrupt innate visual or olfactory preferences. In contrast, loss of *yellow* does influence learning ability, as mutant females failed to modify mate preference following aversive premating experience, indicating an impairment in aversive learning. Despite *yellow*’s role in the melanin biosynthesis pathway, brain dopamine levels remain unchanged in mutant females relative to WT females. These findings identify *yellow* as a pleiotropic gene influencing both pigmentation and aversive mate preference learning, providing evidence that pigmentation genes can shape behavioral processes important for reproductive isolation and speciation.

**Significance statement:** Pigmentation genes are best known for controlling color patterns, but they may also influence behavior through shared neural pathways. Here, we show that the pigmentation gene *yellow* is required for aversive mate preference learning in the butterfly *Bicyclus anynana*. Females lacking *yellow* retain normal innate visual and olfactory mate preferences but fail to learn to avoid previously unattractive mates. Surprisingly, this learning deficit is not associated with altered dopamine levels, suggesting that downstream neural signaling pathways are involved instead. These findings identify a gene that influences both pigmentation and learned mate preference, providing evidence that pigmentation genes can shape behavioral processes important for reproductive isolation and speciation.

## Introduction

Reproductive isolation is a fundamental driver of speciation, where populations diverge genetically and behaviorally to the point of becoming separate species (1, 2). While reproductive barriers can be physical, behavioral, or temporal, maintaining isolation between sympatric and isolated populations often requires the evolution of assortative mate preference and mate choice (3–6). Beyond innate preferences, mate preference learning also plays a role in reproductive isolation by shaping mating behaviors (7). Through social exposure, individuals develop preferences for specific mate traits, often reinforcing reproductive barriers and reducing gene flow (7, 8).

At the genetic level, mutations in pleiotropic genes – where a single gene influences multiple phenotypic traits – can play a significant role in facilitating species divergence. A single pleiotropic gene may simultaneously determine a physical trait important for mate selection while also influencing preference for that trait (5, 9–11). This preference can be innate or learned (7). Some examples of how pleiotropic genes and loci influence both mating trait and innate preference for the trait can be seen in fruitflies (12), Japanese medaka (13), and *Heliconius* butterflies (14–16). However, research on pleiotropic genes that influence both mating traits and learned preferences remains relatively rare. Most genes implicated in mate preference learning are linked to general learning and memory processes, such as neural activity and synaptic plasticity (17–19) like immediate early genes *c-fos* and *zenk* (*egr-1*) (19–23), as well as genes involved in olfaction and vision (24–26). However, although neural and immediate early genes involved in mate preference learning may be pleiotropic within neural systems, they have not been shown to influence both the preferred signal and the learning process shaping preference for that signal. To date, no gene has been demonstrated to couple these two functions. Identifying such pleiotropic genes could therefore provide molecular insights into how mate preference learning may accelerate speciation.

Pigmentation genes involved in melanin production are particularly good candidates for pleiotropically influencing speciation, as pigmentation genes can affect observable traits like camouflage, coloration, visual mating cues, and therefore, influence eventual mate choice (27–29). Insect pigmentation can differ between and within species, and is often a trait important for mate recognition, mate assessment, and mate choice, making it likely to be under sexual selection (27, 28, 30–33). Furthermore, genes controlling wing coloration can also influence pigment production in the eyes of butterflies, altering their visual acuity (34, 35). As a result, pigmentation genes have the potential to impact butterfly vision, their detection of visual mating cues, mate recognition, and the preferred traits themselves, which could in turn drive reproductive isolation and speciation. Recent discoveries have shown that some insect pigmentation genes, such as *tan, ebony,* and *yellow* have pleiotropic roles that extend beyond coloration and may influence neural functions and behavior (36–49). These behavioral effects include altered courtship, aggression, sleep, and vision in pigmentation mutants (especially *ebony* and *tan*) relative to wild-type (WT) individuals. Therefore, pigmentation genes have the potential to facilitate reproductive isolation by simultaneously influencing both color patterns and realized mate preference, either by influencing innate mate preferences, or mate preference learning ability.

One mechanism through which pigmentation genes may influence behavior is via dopamine, a neurotransmitter that is evolutionarily conserved across animals, including cnidarians, nematodes, insects, and vertebrates (50, 51). In vertebrates, dopamine is best known for its roles in arousal, motivation, reward learning, and memory, and disruptions to dopaminergic signaling are implicated in numerous neurological and psychiatric disorders (52–57). In contrast, in arthropods, dopamine is shown to be important in aversive learning instead of reward learning (52, 58–63).

Importantly, dopamine is a key intermediate in the melanin biosynthesis pathway, creating a direct biochemical link between pigmentation genes and behavior via neuromodulatory systems (64, 65). Empirical studies in *Drosophila melanogaster* show that mutations in *ebony* and *tan* pigmentation genes can alter dopamine levels and behavior (36–45, 48), although it remains unknown whether either gene influences preference learning. As for *yellow*, *yellow* mutant *D. melanogaster* exhibited disrupted courtship (46, 47) while *yellow* mutant *Bicyclus anynana* butterflies exhibit increased courtship (49). Unlike *ebony* and *tan*, there are no known studies on the effect of *yellow* on dopamine levels in any insect. Thus, despite dopamine’s central role in either melanin synthesis or aversive learning (66–68), and therefore having the capacity to be influenced by pigmentation genes, there have been no functional studies examining whether pigmentation genes exhibit pleiotropic effects on coloration, dopamine levels, and aversive mate preference learning behavior, and whether this may occur via the dopaminergic pathway.

In this study, we examine the role of the pigmentation gene *yellow* in female innate mate preference and mate preference learning in the butterfly *Bicyclus anynana*. While *yellow* is known to influence melanin pigmentation and courtship intensity in *B. anynana*, its effects on other behaviors remain unexplored (49, 66, 67). Specifically, we investigate its role in innate mate preferences, aversive mate preference learning, and dopamine regulation. To do this, we used CRISPR-Cas9 to knockout *yellow* in *B. anynana*. First, we assessed whether *yellow* mutant females (hereafter *yellow* females) exhibit assortative mating preferences, predicting that they would prefer WT males due to the importance of white UV eyespots in mating preferences, and thus their higher contrast on a darker WT background, compared to a lighter *yellow* background. We then tested whether knocking out *yellow* affected innate visual or olfactory mate preferences. Since *yellow* mutants in *D. melanogaster* have normal visual acuity (41), we expected intact visual discrimination in *yellow* females. Likewise, as *yellow* is not linked to olfactory pathways, we anticipated unaltered olfactory function. Lastly, we examined whether *yellow* influences aversive learning through the dopaminergic pathway. If *yellow* knockout leads to L-DOPA and dopamine accumulation, we hypothesized that *yellow* females would exhibit enhanced aversive learning compared to WT females due to increased dopamine levels. Understanding how *yellow* interacts within the dopaminergic pathway could reveal novel insights into the evolutionary conservation of learning mechanisms across insects by linking pigmentation and learning pathways through shared molecular machinery.

## Results

### Creation of *yellow* mutant line

We generated *yellow B. anynana* mutants using CRISPR-Cas9 by targeting exon 3 of the *yellow* gene (Fig 1A). Different allelic variants were generated in *yellow* mutants: A, B, O (Fig S1B), which originated from some of the WT allelic variants in our population: F, M, S, E (Fig S1A; see also Fig 1B, C). Individuals homozygous for specific alleles tended to be the most common within both our WT and *yellow* female populations (Fig S2). Notably, allele E was not mutated during our injection and line development process, therefore its equivalent is missing from our *yellow* mutants (Fig S2).

**Fig 1:**
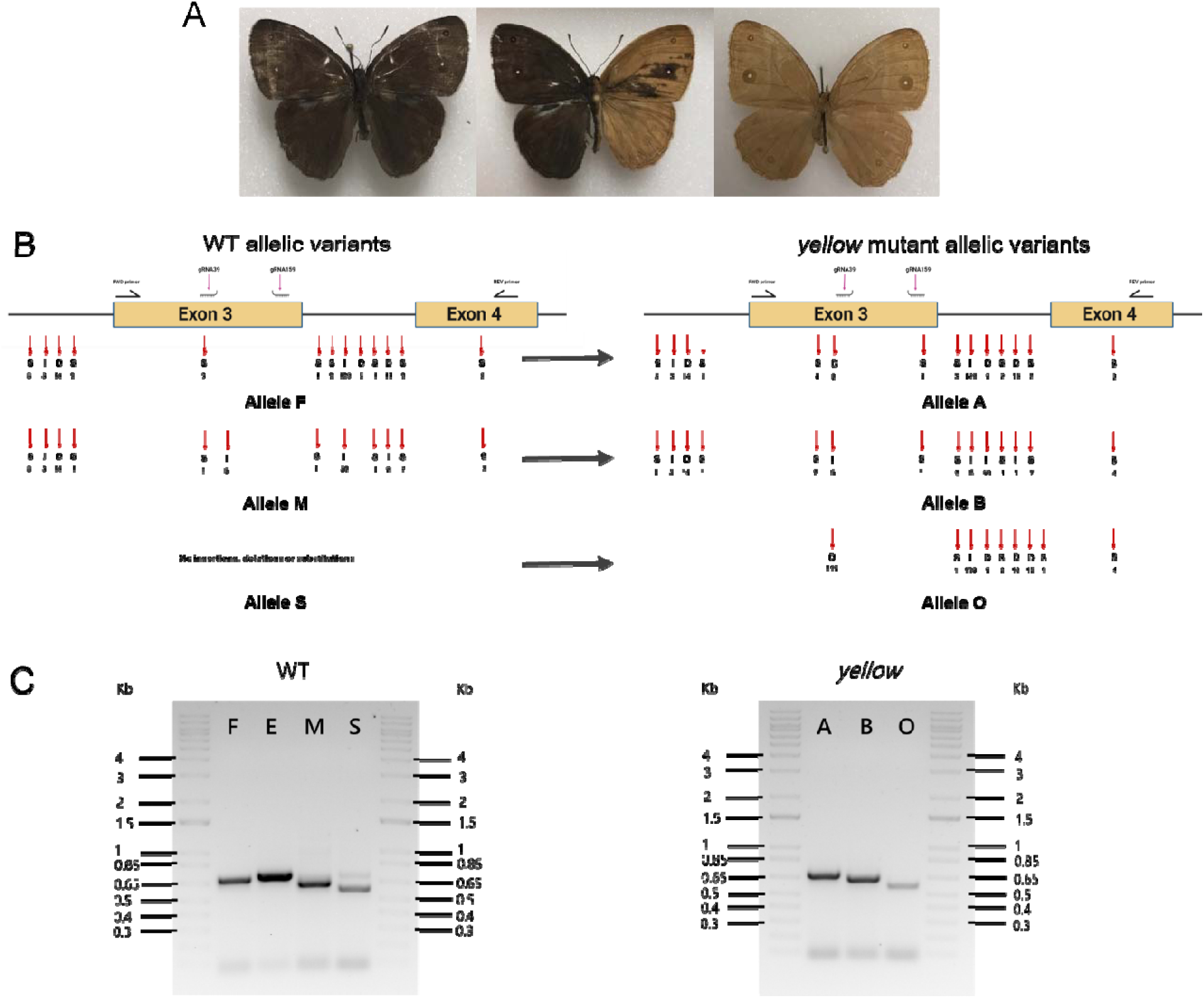
Phenotypes and genotypes of WT and *yellow* mutants. **(A)** Phenotypes of WT, F0 *yellow* mosaic mutant and F2 *yellow* mutants. **(B)** A schematic of the *yellow* gene – Left: The four alleles (F, M, S, and E) that are present in the WT line, and their corresponding nucleotide polymorphisms (E not shown). Right: The three mutant alleles (A, B and O) that are present in the *yellow* line, and their corresponding nucleotide polymorphisms. WT-allele F was mutated to *yellow*-allele A, WT-allele M was mutated to *yellow*-allele B, and WT-allele S was mutated *yellow*-allele O. S, I and D stand for substitutions, insertions and deletions respectively. **(C)** Gel electrophoresis images of the alleles within WT and *yellow* lines.

### *yellow* females mate randomly between *yellow* and WT males

To determine whether *yellow* females had a pre-existing mating bias towards their own phenotype, which have a lighter wing color (Fig. 1A), we conducted mate-choice trials using WT and *yellow* mutant males. We found that *yellow* females mated randomly between *yellow* and WT males (p = 0.655, χ2 = 0.2; Fig. 2A). Therefore, *yellow* females do not mate assortatively by background wing color, and we could standardize male type (WT) for all subsequent comparisons between WT and *yellow* female mate preferences.

**Fig 2.**
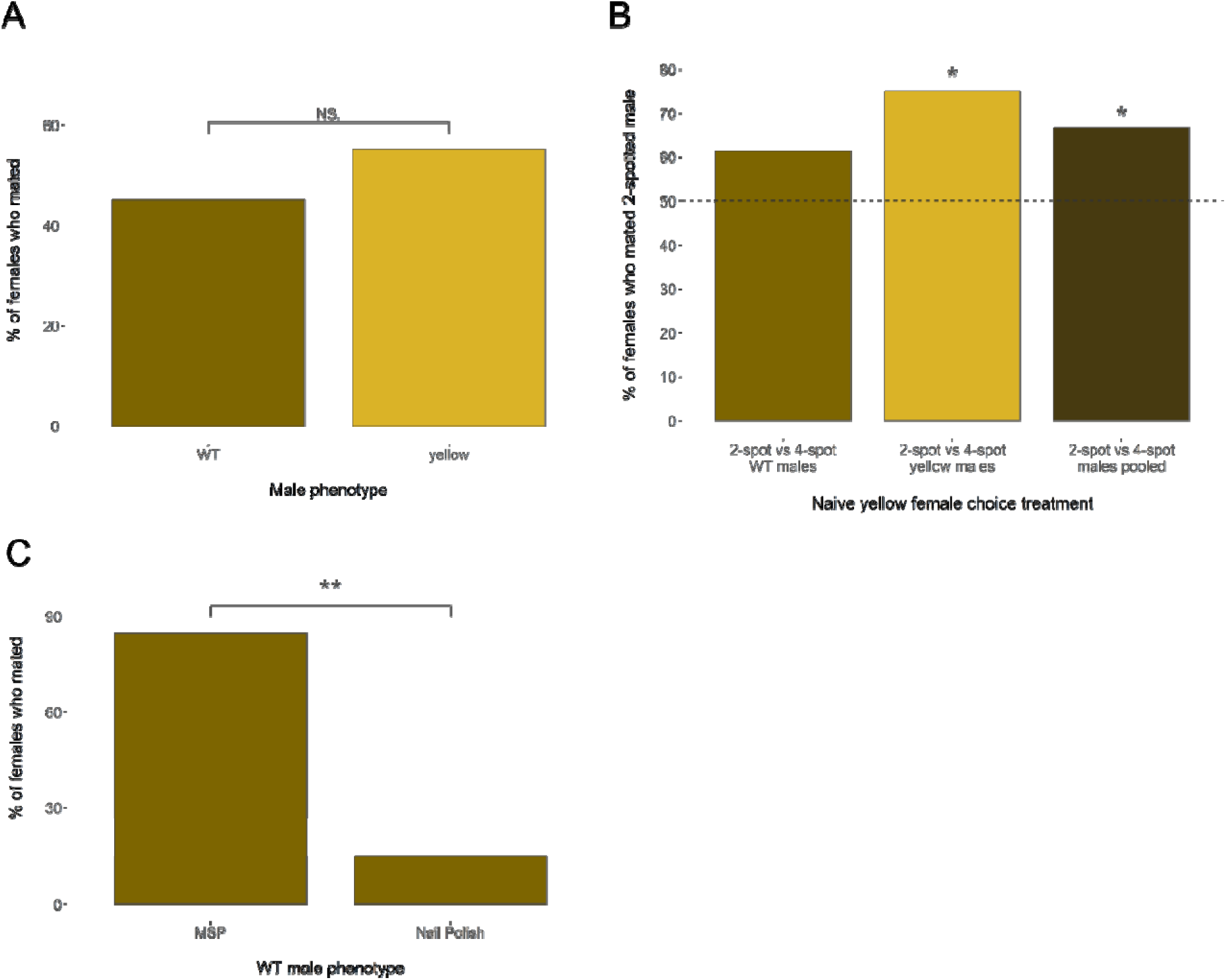
Innate mate preferences in *yellow* females. **(A)** *yellow* females given a choice between *yellow* and WT males mate randomly (n = 20; p = 0.655). **(B)** *yellow* females have an innate preference for 2-spotted males (n = 50; p = 0.0173), though this preference is stronger when choosing between 2 and 4 spotted *yellow* males (n=20; p = 0.0253) than when choosing between 2 and 4 spotted WT males (n = 30; p = 0.209). **(C)** *yellow* females have an innate preference for WT males with undisrupted MSPs over WT males with disrupted pheromone blends (n = 20; p = 0.00174). * = preference significantly different from random. ** = p-value <0.01.

### *yellow* females innately prefer 2-spotted males

Next, we wanted to determine if *yellow* females maintained a pre-existing mating bias towards 2-spotted males, like their WT counterparts (69). We conducted mate-choice trials with both *yellow* males and WT males, giving *yellow* females a choice between 2- and 4-spotted males of each type. Although not statistically significant, *yellow* females tended to prefer mating with 2-spotted over 4-spotted WT males (p = 0.209, χ2 = 1.58; Fig 2B), and had a significant preference for 2-spotted over 4-spotted *yellow* males (p = 0.0253, χ2 = 5; Fig 2B). When we pooled both data sets, we also recovered a significant preference for 2-spotted males (p = 0.0173, χ2 = 5.67; Fig 2B).

This suggests that the *yellow* gene does not influence innate preference for spot number. Therefore, we can use 2-spotted males as our aversive training male wing pattern, as we do for WT females. However, we increased our sample size to n=40 for our learning assays to ensure we would be able to detect effect sizes similar to those previously reported for WT females.

### *Yellow* females dislike WT males with blocked male sex pheromones

We then wanted to determine if *yellow* females maintained a pre-existing mating bias towards males with undisrupted male sex pheromones (MSPs) over males with blocked MSPs, similar to their WT counterparts (70). We conducted mate-choice trials between WT males with undisrupted MSPs and WT males with disrupted MSP blend (i.e. nail polish painted over androconia). We found that *yellow* females preferentially mated with WT males with undisrupted MSP over WT males with disrupted MSP blend (p = 0.00174, χ2 = 9.8; Fig 2D), indicating that the *yellow* mutation did not impact female preference for undisrupted MSPs, and that a disruptive MSP blend is an unpreferred olfactory signal for *yellow* females, just as it is for WT females (70).

### *Yellow* females do not exhibit aversive mate preference learning

Having determined the innate spot and pheromone preferences of *yellow* females, we could then test for an effect of *yellow* on preference learning. To investigate if *yellow* is involved in aversive mate preference learning, we tested for aversive learning ability of *yellow* relative to WT females.

This was done by exposing females (both WT and *yellow*) to 2-spotted WT males with nail polish painted over their androconia for three hours, a treatment that has previously been shown to induce aversive learning in WT females (70), and comparing the mate choice outcomes of these experienced females to those of naïve controls (experienced WT compared to naïve WT, and experienced *yellow* compared to naïve *yellow*). When using naïve *yellow* and WT females as controls, we found that when given a choice between 2-spotted and 4-spotted males, aversively trained WT females avoided mating with innately preferred 2-spotted males and mated preferentially with 4-spotted males compared to naïve WT females, as indicated by the decrease in preference for 2-spotted males (Omnibus p = 0.0394, χ2 = 8.347; pairwise p = 0.0252, χ2 = 5.01; Fig 3), and replicating previously reported results (70). However, trained and naïve *yellow* females did not mate differently from each other (p = 0.361, χ2 = 0.833; Fig 3), suggesting that knocking out *yellow* inhibited the aversive mate preference learning response in this system.

**Fig 3.**
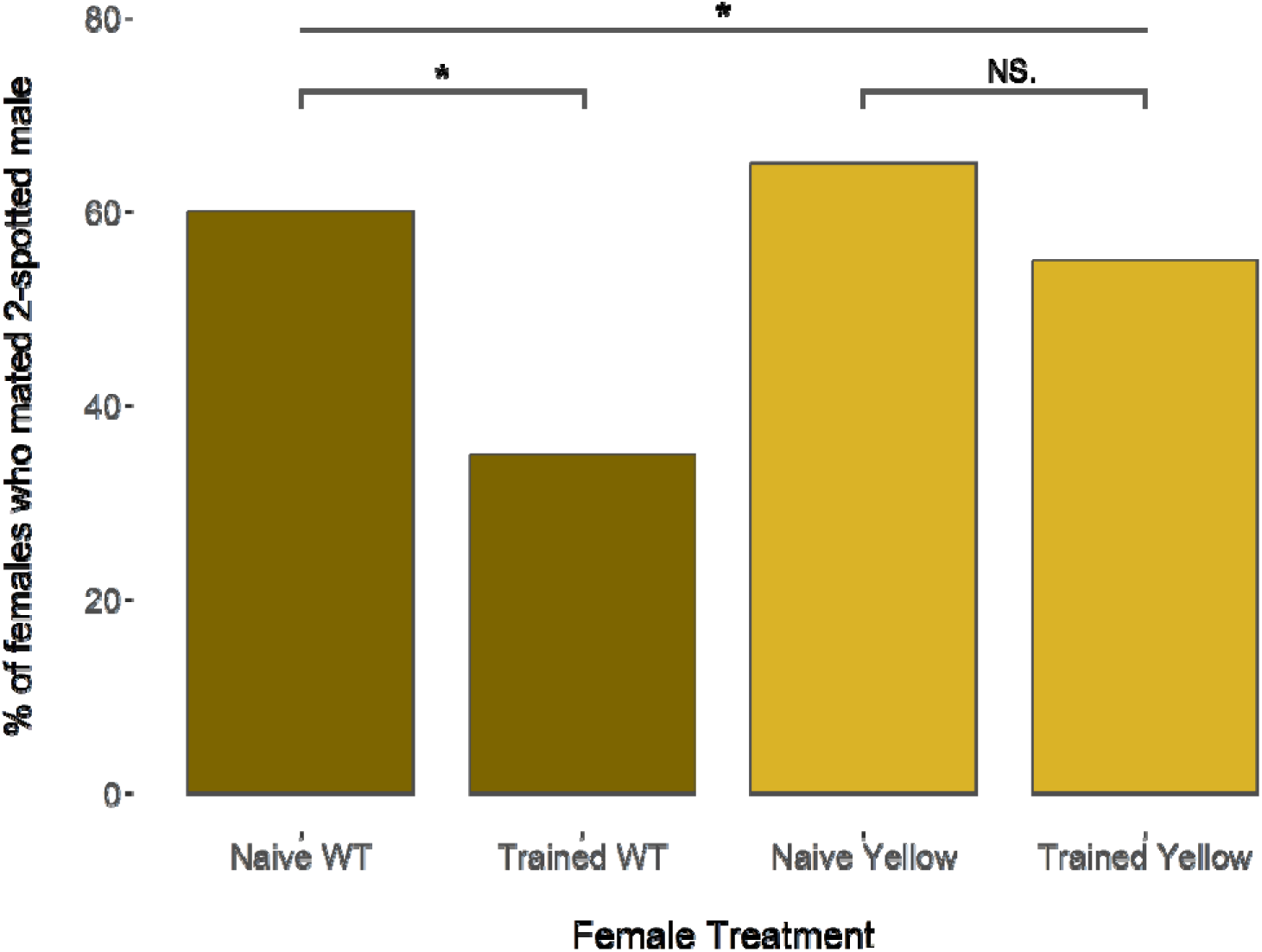
Aversive mate preference learning of WT and *yellow* females. WT females exposed to WT 2-spotted males with a disrupted pheromone blend learn to avoid 2-spotted males and mate more with 4-spotted males (n = 40 per phenotype; p = 0.0252). *Yellow* females exposed to WT 2-spotted males with a disrupted pheromone blend do not learn to avoid 2-spotted males and do not have a preference for 4-spotted males (n = 40 per phenotype; p = 0.361).

### Reduced aversive mate preference learning is not associated with dopamine levels in *B*. anynana females

Next, to investigate if *yellow* influences aversive mate preference learning via the dopaminergic pathway, we quantified dopamine levels using liquid chromatography-mass spectrometry (LC-MS). We found that there was no significant difference between *yellow* and WT females, either when all genotypes within group (*yellow* and WT) were pooled (n = 15 per treatment; t = -0.0472, p = 0. 963), or when individual genotypes within each group were analyzed separately (n = 14 for WT, n = 12 for *yellow;* F = 0.214 and F = 0.774; p = 0.884 and 0.603 respectively) (Fig 4A-C). We also investigated whether these genotypes exhibited variations in pigmented color and UV color, and found that genotypic variation did explain some of the variations in pigmented and UV color within *yellow* and WT lines (Fig S4, S5, Tables S1-S9), even though this genotypic variation did not significantly influence dopamine levels.

**Fig 4.**
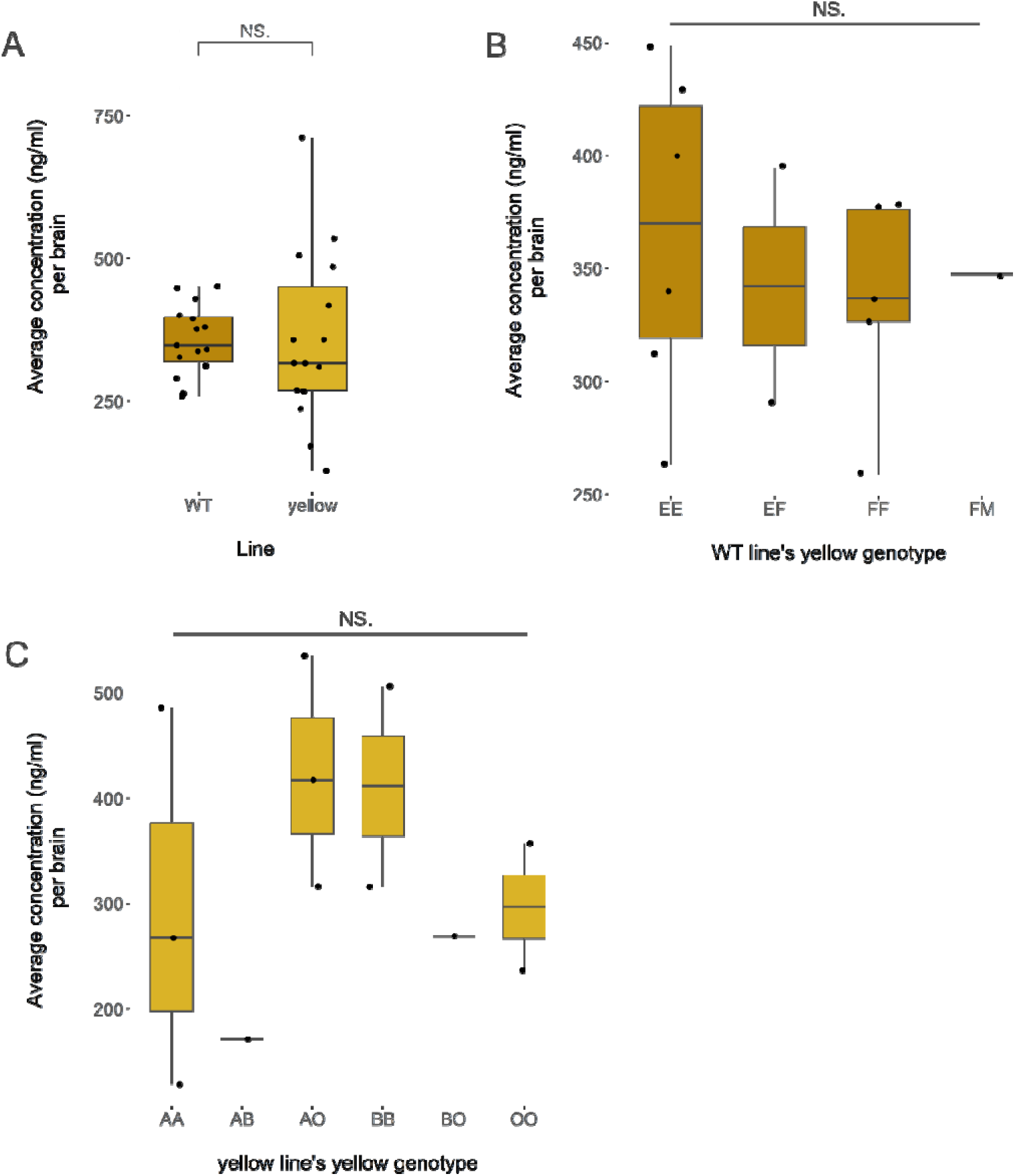
Dopamine levels between WT and *yellow* females. **(A)** Average dopamine concentrations between WT and *yellow* treatments (n = 15 pooled samples per line, p = 0. 963). There were no significant differences between treatments. **(B)** Average dopamine concentrations between *yellow* genotypes within WT females (n = 14 pooled samples; p = 0.884). **(C)** Average dopamine concentrations between *yellow* genotypes within *yellow* females (n = 12 pooled samples ; p = 0.603). For genotype-specific analyses **(B, C)**, pooled samples containing individuals with different genotypes were excluded, resulting in reduced sample sizes. No genotypic variation in dopamine concentrations was detected.

## Discussion

In this study, we showed that the gene *yellow* is necessary for aversive mate preference learning in butterflies, as *yellow* females exposed to 2-spotted males with blocked androconia did not learn to avoid this phenotype in later mating trials the way WT females do. Naïve *yellow* females had similar innate preferences for spot number as WT females, suggesting that the *yellow* mutation did not inhibit female ability to visually distinguish between males with 2 and 4 dorsal forewing spots. Additionally, naive *yellow* females preferred to mate with males with species-specific pheromone blends over males with disrupted pheromones, similar to WT females, suggesting that the *yellow* mutation does not inhibit pheromone detection, or aversion to disrupted pheromones. Thus, the lack of effect of aversive training on *yellow* female’s mating outcome is not due to an inability to detect male signals, but due to an effect of *yellow* on learning and/or memory. This reduction in aversive learning was not mediated by brain dopamine levels in newly emerged females, but may be due to the effect of *yellow* on downstream signaling pathways.

Our finding that *yellow* is a pleiotropic gene affecting both butterfly wing color and mate preference learning of wing patterns, is an illustration of one way a single gene can influence both coloration and mate preference simultaneously. Pleiotropy enables the coupling of signals and preferences, allowing mutations or selection on one trait to drive coordinated evolution of multiple characteristics (71–73). Mutations in pleiotropic genes also introduce variation in both signal and preference, enabling selection to act upon them (72). Most studies on pleiotropic genes (or loci) and mate preference have focused on innate mate preferences (12–15), while research on genes involved in mate preference learning is more limited, though some have been identified in swordtail fishes and butterflies, primarily in neural processing and immediate early genes (17–19). Our study is the first to identify a gene that integrates both coloration and mate preference learning, providing direct evidence that genetic variation in a pigmentation gene can also shape learning processes underlying mate choice. Aversive mate preference learning is shown theoretically to accelerate reproductive isolation (74), as one is able to reinforce preferences by associating con- or heterospecifics with negative experiences (like pheromones). Learned preferences can also become innate through reinforcement by genetic assimilation (75, 76). Therefore, differential aversive mate preference learning abilities within a population can promote behavioral isolation and sympatric speciation when, for example, one learns to avoid a certain trait in mates (i.e. spot number), while the other does not. The intact ability to learn aversively could also lead to better mating decisions and reproductive fitness by reducing hybridization or mating with unfit partners.

The fact that *yellow* mutant females innately exhibited a strong aversion to males with blocked androconia, but did not learn (or remember) to avoid male wing patterns associated with blocked androconia strongly suggests that the genes and/or downstream pathways associated with responding to aversive odor are not the same as the genes associated with transferring negative valence to the visual signal (spot phenotype). This finding aligns with those of studies on the neuroanatomy of associative learning and valence, which show that innate and learned valences are encoded in distinct brain regions in both vertebrates and invertebrates (77, 78). In insects, innate and learned valences are encoded by the lateral horn and mushroom body respectively, with convergence neurons integrating both downstream inputs and regulating behavior (79). The difference does not stop at distinct brain regions, but also molecular pathways, in which different genes are used for innate odor behavior and olfactory learning, mediated by a shared dopaminergic circuit (80). Learned valences, such as exposure to an aversive odor when paired with an innate preferred visual cue (this study), facilitate the switch from innate attraction to learned aversion. Given our finding that knocking out *yellow* inhibits aversion learning, it is possible that *yellow* is involved in the mushroom body’s neural circuitry and not the lateral horn’s, and KO altered connections within the mushroom body. Future studies exploring the neural circuitry and single-cell gene expression of *yellow* and WT female brains may untangle how *yellow* mediates aversive preference learning.

We demonstrated that *yellow* influences aversive learning negatively, but not due to dopamine levels during the developmental time period when training occurs, both of which were contrary to our hypothesis. The unexpected result of impaired aversive learning suggests that dopamine pathways were disrupted, but since dopamine levels remain unchanged, this suggests that downstream dopaminergic pathways were likely affected. Previous studies across multiple insect species have shown that disrupting the dopamine receptors genetically (i.e. RNAi or CRISPR) or pharmacologically (i.e., using antagonists) impair aversive learning (60, 81–84). Therefore, *yellow* mutants may exhibit altered dopamine receptor expression or sensitivity, reducing dopamine’s efficacy in aversive learning. Another potential regulator is 20-hydroxyecdysone (20E), which interacts with dopaminergic signaling through DopEcR (85) and has previously been linked to both aversive memory formation and *yellow* expression (49, 85, 86). Together, these findings suggest that *yellow* may influence aversive learning through broader neuromodulatory or hormonal signaling pathways rather than dopamine levels alone.

Alternatively, *yellow* may indirectly affect learning through processes involved in memory consolidation, such as sleep or circadian rhythm regulation (87). Pigmentation genes such as *ebony* and *tan* have previously been associated with altered sleep patterns (36, 39, 48), raising the possibility that *yellow* may similarly influence circadian processes. Although circadian rhythm was not examined in this study, disrupted sleep cycles could plausibly impair retention of aversive learning experiences in *yellow* females. Future work should therefore investigate dopamine receptor signaling, 20E dynamics, and potential circadian effects in *yellow* mutants.

## Conclusions

Here we show, for the first time in any system, that a pigmentation gene pleiotropically influences both pigmentation and mate preference learning. While *yellow* mutant females retained their innate mate preferences for visual and olfactory cues, they failed to learn aversively, mating indiscriminately with two- and four-spotted males, despite prior exposure to two-spotted males paired with an aversive odor. This suggests that *yellow* is important for aversive mate preference learning, though not through the dopaminergic pathway as initially hypothesized. These findings have implications for reproductive isolation and speciation. If aversive learning reinforces species boundaries by discouraging hybridization, its loss could lead to broader mate choice which hampers reproductive isolation. If learned preferences have a genetic basis, learned preferences can become innate through genetic assimilation and eventually promote speciation when one population learns and one does not. Therefore, mutations in pigmentation genes have the potential to facilitate (or inhibit) reproductive isolation by simultaneously influencing both color and realized mate preference through the mate preference learning pathway.

## Materials and methods

### 1. Study Species and Animal Husbandry

*Bicyclus anynana* is a sub-Saharan, African butterfly that has been reared in the lab since 1988 [71]. The colony at the University of Arkansas was established in 2017, with ∼ 1,000 eggs from a colony at the National University of Singapore. Both WT and *yellow B. anynana* colonies at the University of Arkansas are reared in a climate controlled, USDA-APHIS approved (Permit #526-25-78-75030, # P526-23-44-25151, and Permit # P526P-17-00343) greenhouse. The greenhouse was maintained at 27°C, with a relative humidity of 60-80%, and a 13:11 h light:dark photoperiod provided by 120 V fluorescent lights supplemented with natural light. The larvae were fed with maize plants (*Zea mays*) (Jollytime popcorn) *ad libitum*, and adults were fed with rotten bananas. Adults that emerged on the same day (Day 0) were transferred to sex- and age-specific cages (15.7"L x 15.7"W x 23.6"H).

### 2. Generation of *yellow* mutant line

#### 2.1. sgRNA design and microinjection

The sequence of the *yellow* gene was obtained from the *Bicyclus anynana* v1.2 reference genome (88), scaffold Bany00270. We used the exon 3 sequence to generate sgRNA sequences online using the CRISPOR program (89) (details in Supplemental), specifying the *B. anynana* v1.2 genome and the NGG PAM sequence of Cas9 from *S. pyogenes.* The reverse sgRNA 39 sequence AGCTTTGGAGACGGTTCGTA[GGG], and the forward sgRNA 159 sequence CAGACAAATGTGACCGTCTA[TGG], were selected based on the MIT specificity score of 99 and 100, respectively, thereby minimizing off-target effects in the *B. anynana* genome (PAM sequences in brackets; Fig. S1). Subsequently, the sgRNA sequences were ordered as a Synthetic sgRNA Kit from Synthego (California, USA). Freshly laid eggs were injected with the CRISPR-Cas9 construct mix (300ng/µl of Cas9 - 150ng/µl of gRNA1, and 150ng/µl of gRNA2) using a micromanipulator and syringe (details in Supplemental). Injected eggs were incubated in a mini 20L digital incubator (Ward’s Science; New York, USA) at 27 °C, 13:11 L:D cycle, with fresh corn leaves and balled-up damp Kimwipes to maintain humidity. Upon hatching, individual larvae were transferred to small plastic cups and reared until adulthood. Post-emergence, F0 mosaics were identified, and two mosaic F0 individuals were mated to two WT individuals, producing heterozygous F1s. All F1s were visually WT, confirming that *yellow* KO is recessive for wing color. We then bred F1 individuals, and used the subsequent homozygous *yellow* wing patterned F2 butterflies to build our mutant line.

#### 2.2. PCR amplification and sequencing

PCR amplification reactions to identify *yellow* mutant sequence and WT *yellow* alleles were performed using the forward primer 5’-ATGGTGAGCGCATAGGCATT-3’ and the reverse primer 5’-ACGCGATAAGTCCGTAACCG-3’(IDT; Iowa, USA; Fig. S3). We extracted DNA using a modified chloroform-based method (details in Supplemental). PCR products were analyzed by electrophoresis on a 1% agarose gel in 1X TAE buffer, stained with SYBR Safe (Invitrogen; California, USA), and ran at 100V for 52 minutes. A 1 Kb DNA ladder (Invitrogen; California, USA) was used as a marker. Gels were visualized using a Molecular Imager Gel Doc XR+ with Image Lab 5.0 software (Bio-Rad; California, USA). For sequencing, WT and mutant bands were purified using the QIAquick® PCR Purification Kit (Qiagen; Maryland, USA) and sequenced via Sanger sequencing using the SimpleSeq™ Kit Premixed (Eurofins Genomics; Kentucky, USA). DNA sequence alignments and mutant detection were performed using SnapGene v7.2.

### 3. Behavioral assays

#### 3.1. Testing assortative mating preferences of *yellow* females between *yellow* and WT males

We conducted preliminary assays to determine whether *yellow* females mated assortatively between *yellow* and WT males. For these assays, *yellow* females were kept isolated from morning of emergence from chrysalis (Day 0) to remove effect of social experience on their mate preference (69). On Day 2, one hour after sunrise, we dusted *yellow* females with fluorescent clownfish orange powder (Risk Reactor Inc; California, USA) on their abdomens and then gave them a choice between two virgin, 2-dorsal-forewing-spotted, Day 3 males: one WT and one *yellow* (n = 20). To determine female preference, males were checked using a blacklight flashlight the next day, and the male with orange residue on their abdomen (i.e. mated with the female) was recorded, as orange residue was transferred during copulation. If both males had orange residue, or if no mating occurs after 48 hours, the assay was discarded.

#### 3.2. Testing if *yellow* knockout influenced innate visual spot preference in *yellow* females

Since *yellow* females did not show a strong preference for either *yellow* or WT males (see Results for details), we used WT males for subsequent behavioral assays to control for possible confounding behavioral factors of mutant males. To test if *yellow* knockout affected innate visual spot preference in *yellow* females relative to WT females, we gave naïve Day 2 *yellow* females a choice between 2-spotted and 4-spotted virgin, Day 3 WT males. In previous studies, they showed that WT females innately prefer 2-spotted males (WT) over 4-spotted males (manipulated) (69). To create these male phenotypes, we painted UV-reflective white paint (Reel Wings; North Dakota, USA) on the natural dorsal eyespots of males (2-spotted) or in between the two natural dorsal eyespots of males (4-spotted), as previously described (69). This ensures that both male phenotypes have the same amount of paint, so that female choice is due to number of eyespots rather than presence or amount of paint. Determination of mate choice assays’ outcomes was carried out similar to 3.1 (n = 30).

#### 3.3. Testing effect of male line on spot discrimination in *yellow* females

To test if male *yellow* KO status (i.e., whether the choice males were *yellow* mutants or WT) influenced female spot discrimination, we gave naïve Day 2 *yellow* females a choice between 2-spotted and 4-spotted virgin, Day 3 *yellow* males. Painting of dorsal eyespots in *yellow* males was carried out as mentioned in 3.2. Determination of mate choice assays’ outcomes was carried out similar to 3.1 (n = 20).

#### 3.4. Testing if *yellow* knockout influenced innate olfactory preferences in *yellow* females

Since our subsequent aversive learning assays uses olfactory cues (male sex pheromones; MSP), we tested if the *yellow* knockout affected innate olfactory preferences in *yellow* females. We gave Day 2 *yellow* females a choice between a Day 3, virgin WT male with undisrupted MSPs and a WT male with disrupted MSPs. Previous studies which utilized this experimental design showed that WT females prefer males with undisrupted MSPs over males with disrupted MSPs (70). To disrupt MSPs, we painted Revlon Liquid Quick Dry nail polish (Revlon; New York, USA) over the androconia on the ventral fore- and hindwings of males, as previously described in (70, 90). For males with undisrupted MSPs, we painted nail polish on the anterior part of the dorsal wing of males. This was to make sure that both male phenotypes had nail polish, so that female choice was due to the MSPs being disrupted rather than the presence of the nail polish itself. Determination of mate choice assays’ outcomes was carried out similar to 3.1 (n = 20).

#### 3.5. Aversive learning behavior of *yellow* females

To test if *yellow* knockout influences aversive learning, we exposed newly emerged WT and *yellow* females to sexually mature WT males with disrupted MSPs, and tested the learned response of trained WT and *yellow* females, using naïve WT and *yellow* females as controls. For the aversive learning treatments (e.g. Trained WT or Trained *yellow*), newly emerged Day 0 females were exposed to one Day 3, 2-spotted WT male with disrupted MSPs for 3 hours, after which males were removed and the females kept in isolation, an exposure that has previously been shown to induce an aversive response (70) (n = 40 per treatment). Disruption of MSPs was carried out as mentioned in 3.4. On Day 2, females were given a choice between one 2-spotted and one 4-spotted Day 3 WT. For the naïve treatment (e.g. Naïve WT or Naïve *yellow*), newly emerged Day 0 females were left isolated until mate choice assays, using a similar protocol as that described for the three naïve choice assays described above (n = 40 per treatment). Assays of all treatments were run concurrently, across multiple generations, to control for stochastic variation in greenhouse conditions. Determination of mate choice assays’ outcomes was carried out similar to 3.1.

#### 4. Quantification of dopamine levels in WT and *yellow* females

To quantify whole head dopamine levels, we collected Day 0 females (WT and *yellow*) from August to September 2024 (n = 45 per WT and *yellow* line). They were decapitated at one hour after sunrise, immediately flash frozen in liquid nitrogen and kept at -80°C until dissection.

Genotypes of the *yellow* gene were determined in both WT and *yellow* females. Three heads of the same genotype and treatment were pooled into a tube (n = 15 per treatment) and mechanically lysed in 200ul of 0.1% formic acid (Sigma-Aldrich; Missouri, USA), with 100ng of 3,4-dihydrobenzylamine hydrobromide (3,4-DHBA HBr; Thermo Fisher Scientific; Massachusetts, USA) as an internal standard. Some samples were “mixed” (i.e., had different genotypes pooled) due to not having three heads of the same genotype, these samples were not used when comparing dopamine levels by genotype, but used when comparing dopamine levels by treatment. Details on how samples were prepared and analyzed using LC/MS/MS by the University of Arkansas’ Statewide Mass Spectrometry Facility are in Supplemental Methods.

#### 5. Reflectance spectra measurements

To assess the color patterning in WT and *yellow* mutants, we measured reflectance at seven dorsal and ventral points on the wing (Figs S4, S5) using a Jaz spectrophotometer (Ocean Optics, Inc. Florida, USA). Measurements were performed with the instrument’s internal light source, a flash rate of 200 Hz, and a free-running strobe mode. The probe was positioned at a 45° angle to the wing surface, maintaining a 5 mm distance. Reflectance was recorded from 300 to 700 nm.

### 6. Statistical analyses

#### 6.1. Statistical analyses for behavioral assays

To assess whether *yellow* influences innate mate preferences, we ran goodness-of-fit chi-square analyses to compare the observed females’ choice against an expected random distribution. To assess whether *yellow* influences aversive mate preference learning, we ran chi-square tests of independence to compare between all treatments (omnibus), and between trained and naive treatments in WT and *yellow* females. To assess whether *yellow* influences dopamine levels, we first ran Shapiro-Wilk normality tests and Levene’s tests for equality of variances for each comparison of interest. Since the data were normal and had equal variances, we used ANOVA for all data comparing more than two comparisons (genotype) and independent t-tests for data comparing two groups (treatment). For dopamine quantification analyses, when comparing between genotypes, we removed one WT “mixed” sample (i.e. a mixture of genotypes instead of the same genotypes in one sample), and two *yellow* “mixed” samples and an outlier. Removal of outlier did not affect significance. All analyses were done in R (version 4.4.1).

#### 6.2. Statistical analyses for reflectance spectrometry measurements

To assess differences in color patterning, we calculated the total area under the curve (AUC) for each of the seven points measured in an individual. We performed a one-way analysis of variance (ANOVA) on AUC values using the “stats.f_oneway” function from SciPy. If ANOVA indicated significant differences (p <0.05), we then conducted post-hoc pairwise comparisons using Tukey’s Honestly Significant Difference (HSD) test, using the “pairwise_tukeyhsd” function in Statsmodels. To visualize AUC distribution across genotypes, we generated a boxplot using Seaborn’s “sns.boxplot” function. All statistical analyses for wing patterning were conducted using Python 3.

## 7. Ethical Statement

All *B. anynana* butterflies were maintained in laboratory conditions specified by U.S. Department of Agriculture APHIS permits (Permit #526-25-78-75030, # P526-23-44-25151, and Permit # P526P-17-00343). Butterflies were fed with *ad libitum* food and water. Those used in mate choice assays were sacrificed quickly and frozen in -20°C for future analyses. Those used for dopamine quantification were decapitated quickly and immediately flash frozen, and frozen in -80°C for this and future analyses.

## Acknowledgements

We would like to thank members of the Westerman Lab for their assistance with animal husbandry, and Michael Perry for training in CRISPR/Cas9 gene editing. We would also like to acknowledge the University of Arkansas’ Mass Spectrometry Facility, especially Dr Jennifer Gidden, for their advice and help in running LC-MS. This research was supported by NSF IOS-1937201 and NSF IOS-2243537 to ELW, Animal Behaviour Society’s JEDI Award to YTT, and the University of Arkansas.

## Data availability statement

Behavioral data and codes are available at Dryad (doi.org/10.5061/dryad.7wm37pw7c).

