## Supplemental Material for "The gene *yellow* influences aversive mate preference learning in a butterfly"

**1: Supplemental Figures**

**2: Supplemental Tables**

**3: Detailed Methods**

### Supplementary Figures:

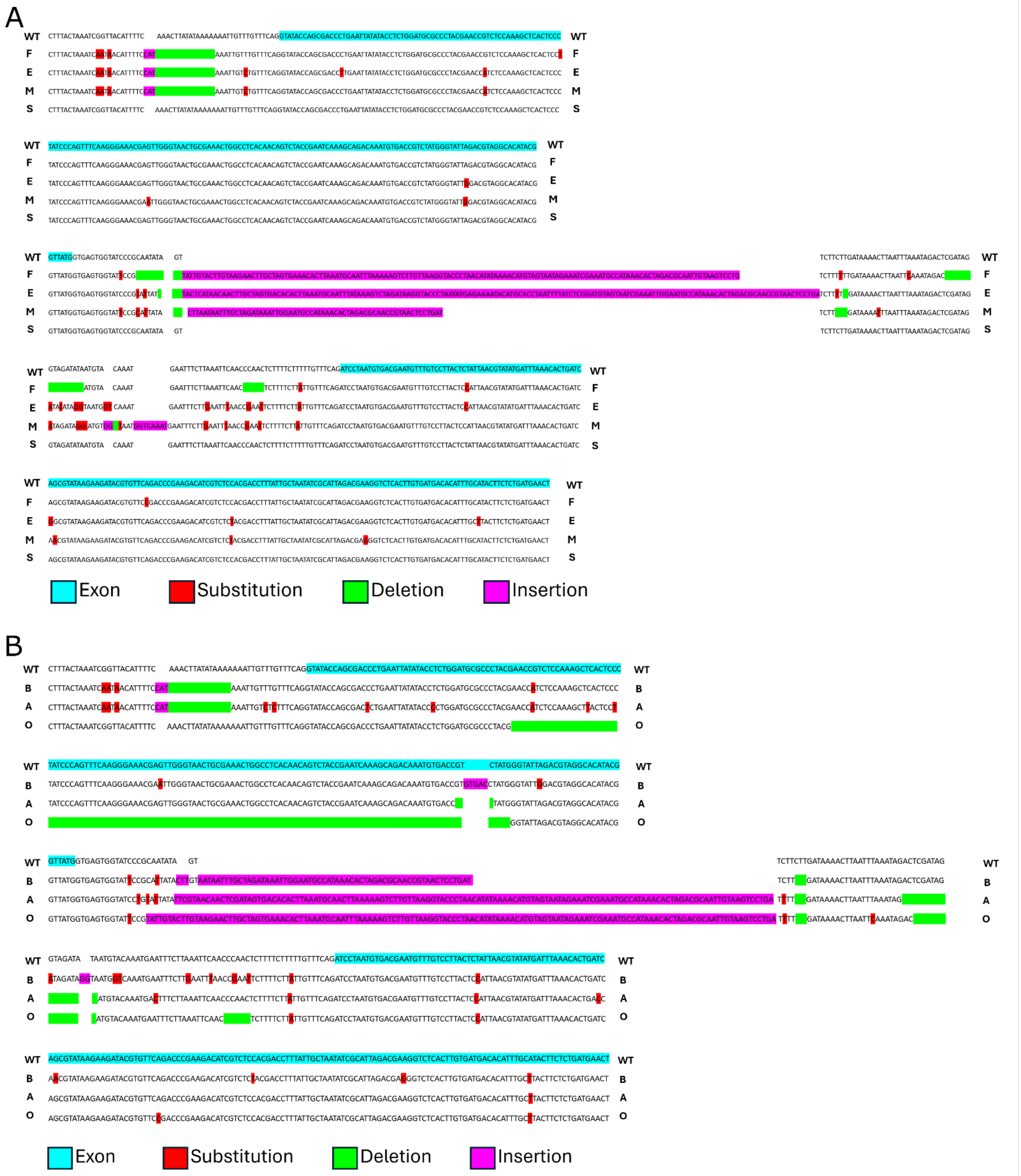

**Fig S1.** Sanger sequencing alignment of wild-type and *yellow* mutant allelic variations. Alignment of genomic DNA sequences comparing the wild-type (WT) reference sequence against various alleles. **(A)** Alignment of WT allelic variations designated as F, E, M, and S. **(B)** Alignment of *yellow* mutant allelic variations designated as B, A, and O. Cyan highlighted blocks indicate exonic regions. Colored indicators denote specific sequence polymorphisms relative to the WT reference: substitutions are shown in red, deletions in green, and insertions in purple.

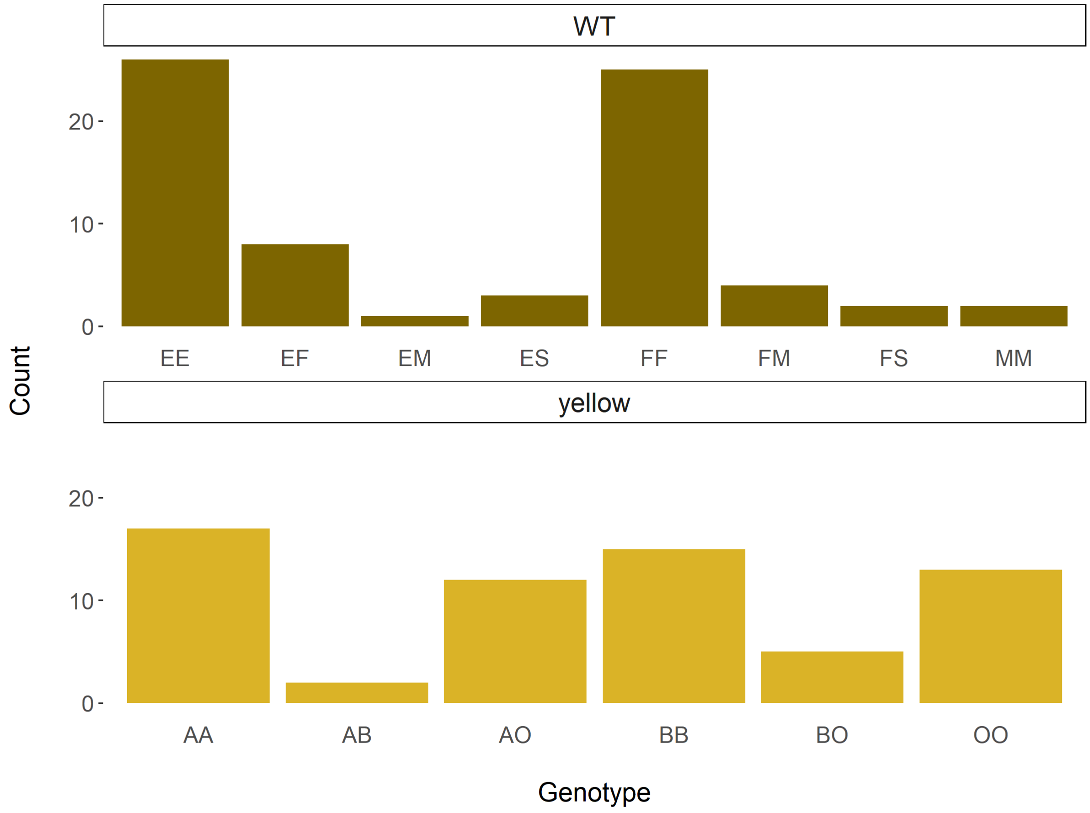


**Fig S2.** Bar graphs of WT and *yellow* genotype frequences within females we collected. Homozygous genotypes tend to be the most common in both populations.

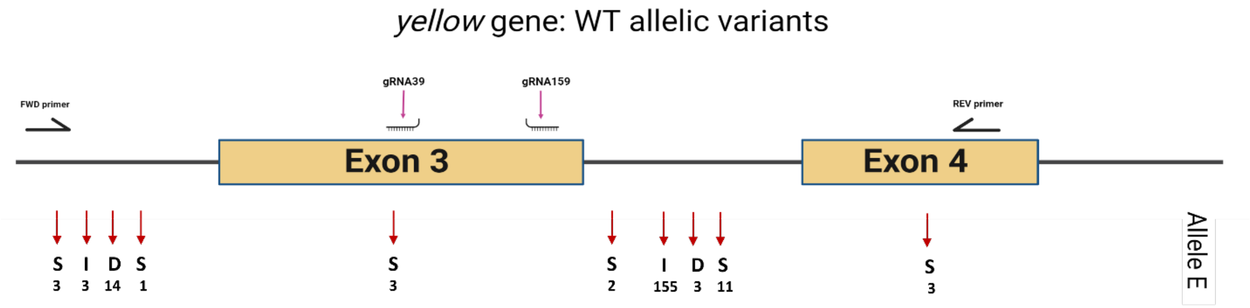


**Fig S3.** Schematic of the *yellow* gene in WT line for allele E, which was not mutated during our line development process, and does not have an equivalent in the *yellow* line.

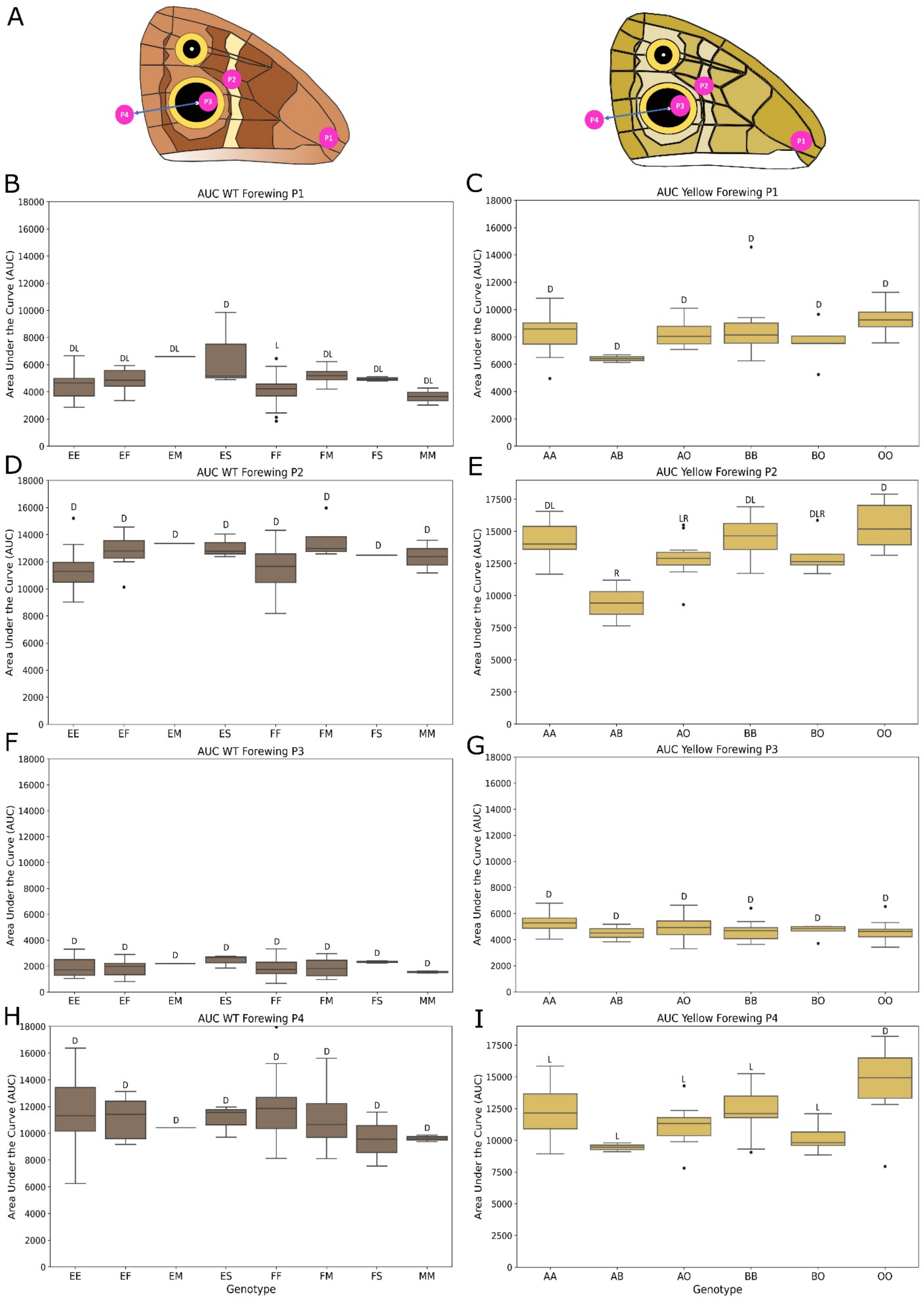
A
P1
P2
P3
P4
B
AUC WT Forewing P1
C
AUC Yellow Forewing P1
D
AUC WT Forewing P2
E
AUC Yellow Forewing P2
F
AUC WT Forewing P3
G
AUC Yellow Forewing P3
H
AUC WT Forewing P4
I
AUC Yellow Forewing P4
Area Under the Curve (AUC)
EE
EF
EM
ES
FF
FM
FS
MM
AA
AB
AO
BB
BO
OO
Genotype

**Fig S4. Boxplots of forewing spectral reflectance for different genotypes in WT & *yellow* females.** **(A)** Schematic of the eight different points measured on the dorsal forewing. **(B, D, F, H)** Measurements of the pigmented scales on WT dorsal forewing, with P4 **(H)** measuring the UV reflective eyespot. **(C,E,G,I)** Measurements of the pigmented scales on *yellow* dorsal forewing, with I representing the UV reflective spot. Genotypes that do not share a letter differ significantly.

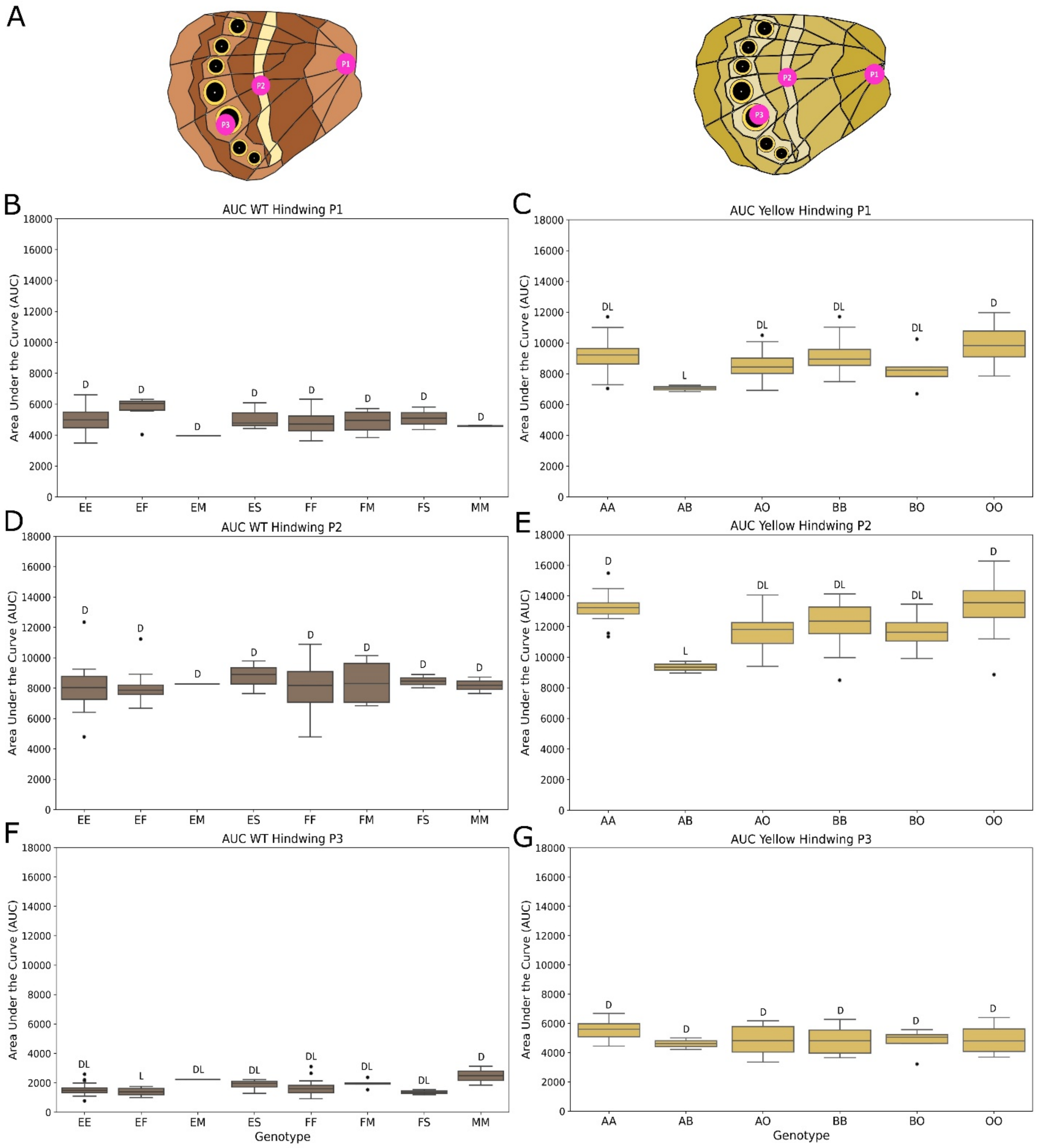


**Fig S5. Boxplots of hindwing spectral reflectance for different genotypes in WT and *yellow* mutant females. (A)** Schematic of the six different points measured on the dorsal hindwing. **(B, D, F)** Measurements of the pigmented scales on the WT dorsal hindwing. **(C,E, G)**

Measurements of the pigmented scales on the *yellow* dorsal hindwing. Genotypes that do not share a letter differ significantly.

#### Supplementary Tables

**Table S1.** Differences between spectral reflectance of WT genotypes at different points of forewing (P1 – P4) and hindwing (P1 – P3). Significant differences are in bold.

| Point | Test | F-statistic | p-value |
|---|---|---|---|
| **Forewing_P1** | **ANOVA** | **3.020185** | **0.008417** |
| **Forewing_P2** | **ANOVA** | **2.654013** | **0.018015** |
| Forewing_P3 | ANOVA | 0.428901 | 0.880568 |
| Forewing_P4 | ANOVA | 0.592247 | 0.759805 |
| Hindwing_P1 | ANOVA | 2.1314 | 0.052874 |
| Hindwing_P2 | ANOVA | 0.143354 | 0.994251 |
| **Hindwing_P3** | **ANOVA** | **2.282957** | **0.038782** |

**Table S2.** Differences between spectral reflectance of *yellow* genotypes at different points of forewing (P1 – P4) and hindwing (P1 – P3). Significant differences are in bold.

| Point | Test | F-statistic | p-value |
|---|---|---|---|
| Forewing_P1 | ANOVA | 2.072846 | 0.0824 |
| **Forewing_P2** | **ANOVA** | **7.029123** | **3.69E-05** |
| Forewing_P3 | ANOVA | 1.959174 | 0.099001 |
| **Forewing_P4** | **ANOVA** | **6.132925** | **0.000134** |
| **Hindwing_P1** | **ANOVA** | **3.453323** | **0.008638** |
| **Hindwing_P2** | **ANOVA** | **4.702145** | **0.001179** |
| Hindwing_P3 | ANOVA | 1.781002 | 0.131668 |

**Table S3.** Pairwise differences of spectral reflectance between WT genotypes at forewing P1. Significant differences are in bold.

| Group1 | Group2 | Mean Diff | Padj | Lower CI | Upper CI |
|---|---|---|---|---|---|
| EE | EF | 345.9631 | 0.9943 | -1073.17 | 1765.1 |
| EE | EM | 2072.493 | 0.6116 | -1504.45 | 5649.435 |
| EE | ES | 2109.714 | 0.0561 | -30.5547 | 4249.983 |
| EE | FF | -414.754 | 0.8869 | -1397.96 | 568.4533 |
| EE | FM | 672.6471 | 0.9503 | -1212.57 | 2557.861 |
| EE | FS | 423.8557 | 0.9995 | -2151.84 | 2999.548 |
| EE | MM | -872.375 | 0.9623 | -3448.07 | 1703.318 |
| EF | EM | 1726.53 | 0.8282 | -1996.47 | 5449.529 |
| EF | ES | 1763.751 | 0.2959 | -612.582 | 4140.084 |
| EF | FF | -760.717 | 0.7044 | -2186.52 | 665.0831 |
| EF | FM | 326.684 | 0.9997 | -1822.79 | 2476.158 |
| EF | FS | 77.8926 | 1 | -2697.07 | 2852.852 |
| EF | MM | -1218.34 | 0.8644 | -3993.3 | 1556.622 |
| EM | ES | 37.2209 | 1 | -4015.87 | 4090.309 |
| EM | FF | -2487.25 | 0.3786 | -6066.84 | 1092.343 |
| EM | FM | -1399.85 | 0.9504 | -5324.23 | 2524.539 |
| EM | FS | -1648.64 | 0.9283 | -5947.59 | 2650.311 |
| EM | MM | -2944.87 | 0.397 | -7243.82 | 1354.081 |
| **ES** | **FF** | **-2524.47** | **0.0105** | **-4669.16** | **-379.776** |
| ES | FM | -1437.07 | 0.6995 | -4117.93 | 1243.799 |
| ES | FS | -1685.86 | 0.7187 | -4890.11 | 1518.389 |
| ES | MM | -2982.09 | 0.0861 | -6186.34 | 222.1582 |
| FF | FM | 1087.401 | 0.6201 | -802.833 | 2977.635 |
| FF | FS | 838.6096 | 0.9698 | -1740.76 | 3417.979 |
| FF | MM | -457.621 | 0.9992 | -3036.99 | 2121.748 |
| FM | FS | -248.791 | 1 | -3288.61 | 2791.025 |
| FM | MM | -1545.02 | 0.7524 | -4584.84 | 1494.794 |
| FS | MM | -1296.23 | 0.9407 | -4806.31 | 2213.846 |

**Table S4.** Pairwise differences of spectral reflectance between WT genotypes at forewing P2. There are no significant differences.

| Group1 | Group2 | Mean Diff | Padj | Lower CI | Upper CI |
|---|---|---|---|---|---|
| EE | EF | 1445.312 | 0.2319 | -396.722 | 3287.346 |
| EE | EM | 2046.826 | 0.8619 | -2596.03 | 6689.681 |
| EE | ES | 1766.135 | 0.4947 | -1011.92 | 4544.195 |
| EE | FF | 128.3873 | 1 | -1147.81 | 1404.586 |
| EE | FM | 2325.37 | 0.0742 | -121.63 | 4772.369 |
| EE | FS | 1190.074 | 0.9509 | -2153.16 | 4533.312 |
| EE | MM | 1072.998 | 0.9718 | -2270.24 | 4416.235 |
| EF | EM | 601.5145 | 0.9999 | -4230.92 | 5433.951 |
| EF | ES | 320.823 | 1 | -2763.65 | 3405.293 |
| EF | FF | -1316.92 | 0.3481 | -3167.61 | 533.7575 |
| EF | FM | 880.0576 | 0.9744 | -1909.95 | 3670.066 |
| EF | FS | -255.238 | 1 | -3857.12 | 3346.648 |
| EF | MM | -372.314 | 1 | -3974.2 | 3229.571 |
| EM | ES | -280.692 | 1 | -5541.58 | 4980.199 |
| EM | FF | -1918.44 | 0.8976 | -6564.73 | 2727.853 |
| EM | FM | 278.5431 | 1 | -4815.29 | 5372.378 |
| EM | FS | -856.752 | 0.9997 | -6436.77 | 4723.264 |
| EM | MM | -973.829 | 0.9993 | -6553.85 | 4606.188 |
| ES | FF | -1637.75 | 0.5931 | -4421.55 | 1146.054 |
| ES | FM | 559.2346 | 0.9996 | -2920.52 | 4038.986 |
| ES | FS | -576.061 | 0.9999 | -4735.16 | 3583.038 |
| ES | MM | -693.137 | 0.9995 | -4852.24 | 3465.962 |
| FF | FM | 2196.982 | 0.1116 | -256.534 | 4650.498 |
| FF | FS | 1061.687 | 0.9737 | -2286.32 | 4409.697 |
| FF | MM | 944.6105 | 0.9865 | -2403.4 | 4292.62 |
| FM | FS | -1135.3 | 0.9848 | -5080.96 | 2810.372 |
| FM | MM | -1252.37 | 0.9735 | -5198.04 | 2693.296 |
| FS | MM | -117.076 | 1 | -4673.14 | 4438.988 |

**Table S5.** Pairwise differences of spectral reflectance between WT genotypes at hindwing P3. Significant differences are in bold.

| Group1 | Group2 | Mean Diff | Padj | Lower CI | Upper CI |
|---|---|---|---|---|---|
| EE | EF | -179.074 | 0.9724 | -738.903 | 380.7546 |
| EE | EM | 662.6461 | 0.819 | -748.405 | 2073.697 |
| EE | ES | 294.9062 | 0.9556 | -549.398 | 1139.211 |
| EE | FF | 72.2397 | 0.999 | -315.621 | 460.1005 |
| EE | FM | 387.4592 | 0.7286 | -356.23 | 1131.148 |
| EE | FS | -192.067 | 0.9989 | -1208.14 | 824.0057 |
| EE | MM | 921.5473 | 0.1026 | -94.5253 | 1937.62 |
| EF | EM | 841.7203 | 0.6246 | -626.948 | 2310.389 |
| EF | ES | 473.9804 | 0.7572 | -463.448 | 1411.409 |
| EF | FF | 251.3139 | 0.8535 | -311.143 | 813.771 |
| EF | FM | 566.5334 | 0.4296 | -281.403 | 1414.469 |
| EF | FS | -12.9928 | 1 | -1107.67 | 1081.688 |
| **EF** | **MM** | **1100.622** | **0.0478** | **5.9409** | **2195.302** |
| EM | ES | -367.74 | 0.996 | -1966.62 | 1231.143 |
| EM | FF | -590.406 | 0.8914 | -2002.5 | 821.6891 |
| EM | FM | -275.187 | 0.9992 | -1823.3 | 1272.925 |
| EM | FS | -854.713 | 0.7602 | -2550.59 | 841.1588 |
| EM | MM | 258.9012 | 0.9997 | -1436.97 | 1954.773 |
| ES | FF | -222.667 | 0.991 | -1068.72 | 623.3831 |
| ES | FM | 92.553 | 1 | -965.009 | 1150.115 |
| ES | FS | -486.973 | 0.9267 | -1751 | 777.0551 |
| ES | MM | 626.6411 | 0.7749 | -637.387 | 1890.669 |
| FF | FM | 315.2195 | 0.8857 | -430.45 | 1060.889 |
| FF | FS | -264.307 | 0.9917 | -1281.83 | 753.2165 |
| FF | MM | 849.3076 | 0.1694 | -168.216 | 1866.831 |
| FM | FS | -579.526 | 0.7965 | -1778.69 | 619.6364 |
| FM | MM | 534.0881 | 0.8556 | -665.074 | 1733.251 |
| FS | MM | 1113.614 | 0.2057 | -271.059 | 2498.288 |

**Table S6.** Pairwise differences of spectral reflectance between *yellow* genotypes at forewing P2. Significant differences are in bold.

| Group1 | Group2 | Mean Diff | Padj | Lower CI | Upper CI |
|---|---|---|---|---|---|
| **AA** | **AB** | **-4878.78** | **0.0013** | **-8307.31** | **-1450.25** |
| AA | AO | -1383.6 | 0.1879 | -3112.84 | 345.641 |
| AA | BB | 212.1519 | 0.999 | -1477.65 | 1901.953 |
| AA | BO | -1136.39 | 0.7046 | -3469.71 | 1196.918 |
| AA | OO | 955.8711 | 0.5575 | -733.93 | 2645.672 |
| AB | AO | 3495.181 | 0.0508 | -7.7329 | 6998.095 |
| **AB** | **BB** | **5090.931** | **0.0009** | **1607.317** | **8574.545** |
| AB | BO | 3742.385 | 0.0599 | -94.8654 | 7579.635 |
| **AB** | **OO** | **5834.65** | **0.0001** | **2351.036** | **9318.264** |
| AO | BB | 1595.75 | 0.1234 | -240.276 | 3431.776 |
| AO | BO | 247.2035 | 0.9997 | -2194.09 | 2688.497 |
| **AO** | **OO** | **2339.469** | **0.0052** | **503.4432** | **4175.495** |
| BB | BO | -1348.55 | 0.5706 | -3762.07 | 1064.972 |
| BB | OO | 743.7192 | 0.8253 | -1055.21 | 2542.65 |
| BO | OO | 2092.266 | 0.1252 | -321.253 | 4505.784 |

**Table S7.** Pairwise differences of spectral reflectance between *yellow* genotypes at forewing P4 (UV eyespot). Significant differences are in bold.

| Group1 | Group2 | Mean Diff | Padj | Lower CI | Upper CI |
|---|---|---|---|---|---|
| AA | AB | -2864.09 | 0.4207 | -7342.75 | 1614.567 |
| AA | AO | -1174.69 | 0.6439 | -3433.57 | 1084.201 |
| AA | BB | -50.8934 | 1 | -2258.26 | 2156.476 |
| AA | BO | -2112.18 | 0.331 | -5160.16 | 935.802 |
| **AA** | **OO** | **2339.018** | **0.0318** | **131.6485** | **4546.388** |
| AB | AO | 1689.406 | 0.8836 | -2886.41 | 6265.227 |
| AB | BB | 2813.199 | 0.4591 | -1737.41 | 7363.808 |
| AB | BO | 751.9113 | 0.9977 | -4260.65 | 5764.472 |
| **AB** | **OO** | **5203.11** | **0.0162** | **652.5008** | **9753.72** |
| AO | BB | 1123.793 | 0.7371 | -1274.59 | 3522.174 |
| AO | BO | -937.495 | 0.9527 | -4126.53 | 2251.542 |
| **AO** | **OO** | **3513.704** | **0.0009** | **1115.323** | **5912.086** |
| BB | BO | -2061.29 | 0.3957 | -5214.04 | 1091.468 |
| **BB** | **OO** | **2389.912** | **0.044** | **39.9871** | **4739.836** |
| **BO** | **OO** | **4451.199** | **0.0015** | **1298.444** | **7603.954** |

**Table S8.** Pairwise differences of spectral reflectance between *yellow* genotypes at hindwing P1. Significant differences are in bold.

| Group1 | Group2 | Mean Diff | Padj | Lower CI | Upper CI |
|---|---|---|---|---|---|
| AA | AB | -2057.89 | 0.1758 | -4593.65 | 477.8739 |
| AA | AO | -505.648 | 0.8506 | -1784.6 | 773.3059 |
| AA | BB | 160.6597 | 0.9989 | -1089.13 | 1410.446 |
| AA | BO | -821.73 | 0.7239 | -2547.46 | 904.0003 |
| AA | OO | 767.1283 | 0.4671 | -482.658 | 2016.914 |
| AB | AO | 1552.24 | 0.4943 | -1038.53 | 4143.013 |
| AB | BB | 2218.548 | 0.1299 | -357.952 | 4795.047 |
| AB | BO | 1236.158 | 0.7918 | -1601.89 | 4074.208 |
| **AB** | **OO** | **2825.016** | **0.0238** | **248.5167** | **5401.516** |
| AO | BB | 666.3081 | 0.698 | -691.626 | 2024.243 |
| AO | BO | -316.082 | 0.9953 | -2121.68 | 1489.512 |
| AO | OO | 1272.777 | 0.0784 | -85.1578 | 2630.711 |
| BB | BO | -982.39 | 0.5866 | -2767.44 | 802.6612 |
| BB | OO | 606.4686 | 0.7588 | -724.03 | 1936.967 |
| BO | OO | 1588.859 | 0.1078 | -196.193 | 3373.91 |

**Table S9.** Pairwise differences of spectral reflectance between *yellow* genotypes at hindwing P2. Significant differences are in bold.

| Group1 | Group2 | Mean Diff | Padj | Lower CI | Upper CI |
|---|---|---|---|---|---|
| **AA** | **AB** | **-3864.6** | **0.0096** | **-7070.54** | **-658.664** |
| AA | AO | -1519.07 | 0.0772 | -3136.04 | 97.8978 |
| AA | BB | -942.48 | 0.4993 | -2522.57 | 637.611 |
| AA | BO | -1553.57 | 0.3017 | -3735.39 | 628.2507 |
| AA | OO | 125.7207 | 0.9999 | -1454.37 | 1705.811 |
| AB | AO | 2345.53 | 0.2957 | -929.957 | 5621.017 |
| AB | BB | 2922.12 | 0.1029 | -335.32 | 6179.561 |
| AB | BO | 2311.028 | 0.4127 | -1277.09 | 5899.145 |
| **AB** | **OO** | **3990.321** | **0.0081** | **732.8804** | **7247.761** |
| AO | BB | 576.5905 | 0.9189 | -1140.23 | 2293.412 |
| AO | BO | -34.5014 | 1 | -2317.29 | 2248.291 |
| AO | OO | 1644.791 | 0.0678 | -72.0309 | 3361.613 |
| BB | BO | -611.092 | 0.9665 | -2867.91 | 1645.729 |
| BB | OO | 1068.2 | 0.4287 | -613.935 | 2750.335 |
| BO | OO | 1679.292 | 0.2562 | -577.529 | 3936.113 |

**Detailed methods:**

1. **Study Species and Animal Husbandry**

*Bicyclus anynana* is a sub-Saharan, African butterfly that has been reared in the lab since 1988 (1). The colony at the University of Arkansas was established in 2017, with ~ 1,000 eggs from a colony at the National University of Singapore. Both WT and *yellow B. anynana* colonies at the University of Arkansas are reared in a climate controlled, USDA-APHIS approved (Permit #526-25-78-75030, # P526-23-44-25151, and Permit # P526P-17-00343) greenhouse. The greenhouse was maintained at 27ºC, with a relative humidity of 60-80%, and a 13:11 h light:dark photoperiod provided by 120 V fluorescent lights supplemented with natural light. The larvae were fed with maize plants (*Zea mays*) (Jollytime popcorn) *ad libitum*, and adults were fed with rotten bananas. Adults that emerged on the same day (Day 0) were transferred to sex- and age-specific cages (15.7"L x 15.7"W x 23.6"H).

2. **Generation of *yellow* mutant line**

**2.1. sgRNA Design and microinjection**

The sequence of the *yellow* gene was obtained from the *Bicyclus anynana* v1.2 reference genome (2), scaffold Bany00270. We used the exon 3 sequence to generate sgRNA sequences online using the CRISPOR program (3), specifying the *B. anynana* v1.2 genome and the NGG PAM sequence of Cas9 from *S. pyogenes.* The reverse sgRNA 39 sequence AGCTTTGGAGACGGTTCGTA[GGG], and the forward sgRNA 159 sequence CAGACAAATGTGACCGTCTA[TGG], were selected based on the MIT specificity score of 100 and 99, respectively, thereby minimizing off-target effects in the *B. anynana* genome (PAM sequences in brackets; Fig. S1). Subsequently, the sgRNA sequences were ordered as a Synthetic sgRNA Kit from Synthego (California, USA).

Fresh *Zea mays* plants were placed in the WT colony cages for a 1.5-hour egg collection period. The collected eggs were washed with 7.5% benzalkonium chloride for 30 seconds, followed by three washing steps with ultrapure water, each lasting 1 minute. The eggs were then dried for 10 minutes and attached to microscope slides using double-sided tape strips with a fine wet brush. For the injection process, we used a micromanipulator MN-153 attached to a magnetic stand GJ-1 and an iron plate IP (Narishige; New York, USA), along with a borosilicate needle, prepared with a PUL-1000 (World Precision Instruments; Florida, USA). The needle was connected to a 50 mL syringe with Tygon E-3603 tubing (1/32 x 3/32") to permit injection. Then, the CRISPR-Cas9 construct mix (300ng/µl of Cas9 - 150 ng/ul of gRNA1, and 150 ng/ul of gRNA2) was loaded into the needle by pulling up on the plunger of the attached syringe. During the injection, the needle was positioned at a 45° angle, the eggs were punctured until yolk flowed into or around the needle at the egg's surface, and a small volume of construct was injected. Injected eggs were incubated in a mini 20L digital incubator (Ward's Science; New York, USA) at 27 °C, with fresh corn leaves and balled-up damp Kimwipes to maintain humidity. Upon hatching, individual larvae were transferred to small plastic cups and reared until adulthood. Post-emergence, F0 mosaics were identified, and two mosaic F0 individuals were mated to two WT individuals, producing heterozygous F1s. All F1s were visually WT, confirming that *yellow* KO is recessive for wing color. We then bred F1 individuals, and used the subsequent homozygous *yellow* wing patterned F2 butterflies to build our mutant line.

##### 2.2. PCR amplification and sequencing

PCR amplification reactions to identify *yellow* mutant sequence and WT *yellow* alleles were performed using the forward primer 5'-ATGGTGAGCGCATAGGCATT-3' and the reverse primer 5'-ACGCGATAAGTCCGTAACCG-3'(IDT; Iowa, USA; Fig. S3). We extracted DNA using a modified chloroform-based method, where leg tissues were homogenized in 250µl extraction buffer supplemented with 3µl Proteinase K, 5µl 2-mercaptoethanol and 1µl RNase. PCR reactions were prepared in a 20µl volume containing 9.6µl nuclease-free water, 4µl of 5x Green GoTaq Reaction Buffer, 2µl of 2mM dNTPs mix, 1µl of 10 µM forward primer, 1µl of 10 µM reverse primer, 0.4µl of Taq Polymerase, and 2µl of DNA template. Amplifications were performed using a T100™ Thermal Cycler (BioRad;

California, USA) under the following conditions: initial denaturation at 94°C for 4 minutes, followed by 35 cycles of 30 seconds at 94°C (denaturation), 45 seconds at 60°C (annealing), 36 seconds at 72°C (extension), with a final extension at 72°C for 5 minutes. PCR products were analyzed by electrophoresis on a 1% agarose gel in 1X TAE buffer, stained with SYBR Safe, and ran at 100V for 52 minutes. A 1 Kb DNA ladder was used as a marker. Gels were visualized using a Molecular Imager Gel Doc XR+ with Image Lab 5.0 software (Bio-Rad; California, USA). For sequencing, WT and mutant bands were purified using the QIAquick® PCR Purification Kit (Qiagen; Maryland, USA) and sequenced via Sanger sequencing using the SimpleSeq™ Kit Premixed (Eurofins Genomics; Kentucky, USA). DNA sequence alignments and mutant detection were performed using SnapGene v7.2.

(Sigma-Aldrich; Missouri, USA), with 100ng of 3,4-dihydrobenzylamine hydrobromide (3,4-DHBA HBr; Thermo Fisher Scientific; Massachusetts, USA) as an internal standard. Some samples were "mixed" (i.e. had different genotypes pooled) due to not having three heads of the same genotype, these samples were not used when comparing dopamine levels by genotype, but used when comparing dopamine levels by treatment. Samples were centrifuged at 17,100 x g for 20 minutes at 4°C, twice more for 5 minutes for sufficient homogenization, and solution of 1:10 n-hexane (Sigma-Aldrich; Missouri, USA): ultrapure water was added to the supernatant. The samples were centrifuged at 17,100 x g for 5 minutes at 4°C. 100 ul of the aqueous (bottom) layer was extracted into 250 ul inserts (Restek; Pennsylvania, USA) in 2 ml, 9mm amber autosampler vials (VWR; Pennsylvania, USA), and sent to the University of Arkansas Statewide Mass Spectrometry Facility for LC/MS/MS analysis. 2ul of each sample was injected into Shimadzu LCMS-8040 (Maryland, USA), and separated by a Kinetex Polar C18 column (2.6um, 100 x 2.1mm; Phenomenex, California, USA) at 40°C, using an isocratic run at 2% CAN and 98% water with 0.1% formic acid added as an eluent solution at 0.2 ml/min flow rate. Multiple reaction monitoring (MRM) transitions for each compound (parent m/z → daughter m/z) were applied as follows: dopamine, 154.0 → 137.0; and DHBA, 140.0 → 123.0. The retention time of dopamine was 2.48 min, and the retention time of DHBA was 2.25 min. The amounts of dopamine and DHBA were calculated using calibration curves of respective external standards with 100ng of DHBA.
